# Fingerprints as Frozen Nematic Fields

**DOI:** 10.64898/2025.12.19.695577

**Authors:** Lorna Merrett, Kimia Witte

## Abstract

Human fingerprints arise during embryogenesis and persist throughout life. While biochemical mechanisms of ridge initiation and growth are increasingly well understood, the physical principles governing ridge-pattern organisation remain unclear. Here, we show that fingerprint ridges can be treated as a frozen nematic field on a bounded surface, exhibiting long-range orientational order and singularities. Modelling ridge orientation as a two-dimensional nematic director field on the functional contact area of the fingertip, we apply a generalised Poincaré–Hopf theorem to derive a topological constraint on permitted configurations. Triradii, loops, and whorls correspond to 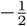, 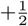, and +1 nematic defects, whose total charge is set by surface topology and boundary winding. Across 133 fingerprints from open-access databases, 95.5% satisfy the predicted charge-neutrality condition. Although boundary winding is typically below the idealised value (*W*_∂*S*_ = 0.71 ± 0.12 vs. *W*_∂*S*_ ≈1), fingerprints above a threshold (*W*_∂*S*_ ≥ 0.6) consistently maintain zero net charge, revealing a coarse-grained realisation of the underlying topological rule. These results identify a mathematical limit on allowable fingerprint configurations and position fingerprints as a passive biological record of nematic order, with topology acting as a fundamental constraint on developmental pattern formation.

## 2 Introduction

Nematic liquid crystals have two defining features: long-range orientational order and the presence of topological defects [1, 2]. These defects act as reliable organising centres in morphogenesis, and growth and repair fail when they have been disrupted [3, 4]. *In vitro*, they serve as mechanotransduction sites where extrusion and extension into the third dimension occur [5, 6]. Long-range nematic order has also been shown to help maintain structure during early embryogenesis across many species [7]. Biology, therefore, appears to make extensive use of the characteristic properties of the nematic phase in developmental processes and self-organisation across scales.

The human fingerprint, studied in the field of dermatoglyphics, naturally embodies nematic features: ridges are strongly aligned in large regions and distinct patterns disrupt this order. The literature conventionally focuses on the generative mechanisms of fingerprints, and the spread of ridges has recently been shown to be initiated by biochemical signalling from key sites on the finger pad [8]. In contrast, mathematical and conceptual approaches characterise the hand as a topological tensor field [9, 10], linking the number of digits with the number of patterns across the fingers and palm.

In this view, the fingerprint becomes a static record of orientational order on a confined domain, constrained by geometry rather than driven by activity. Using this nematic framework, we do not claim that fingerprint ridges constitute an active nematic in the strict biophysical sense during embryogenesis. Instead, we propose that the mathematical framework of nematics provides an intuitive language for describing ridge orientation fields and their topological constraints, widening the application of nematics in biology. This approach complements existing models: biochemical pathways initiate and propagate ridges [8], while a mathematical approach treats patterns as singularities in a continuous global field [9]. A nematic perspective provides geometric rules, local to the fingertip, that connect topology to permitted pattern configurations, bridging the conceptual and the physical.

This work presents a simple application of nematics to human fingerprints, treating the ridges as a frozen nematic field on a surface to explain why certain configurations are observed across almost all human fingers. By neglecting complex signalling and mechanical intricacies, we adopt a local, nematic-based topological defect perspective that predicts a conserved net charge of zero for individual fingerprints, revealing a direct relationship between topology and biology.

## 3 Fingerprint patterns as nematic defects

The patterns that arise in fingerprints are strongly reminiscent of topological defects in nematic systems. Triradii, loops, and whorls correspond to 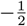, 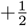, and +1 defects respectively (shown in Fig. 1), where the defect charge describes the winding (rotation) of the director field around a singular point [2, 9]. These defect types are readily identified and form the basis of classical fingerprint classification. Despite this, fingerprints have not previously been analysed explicitly within a nematic framework.

**Figure 1.**
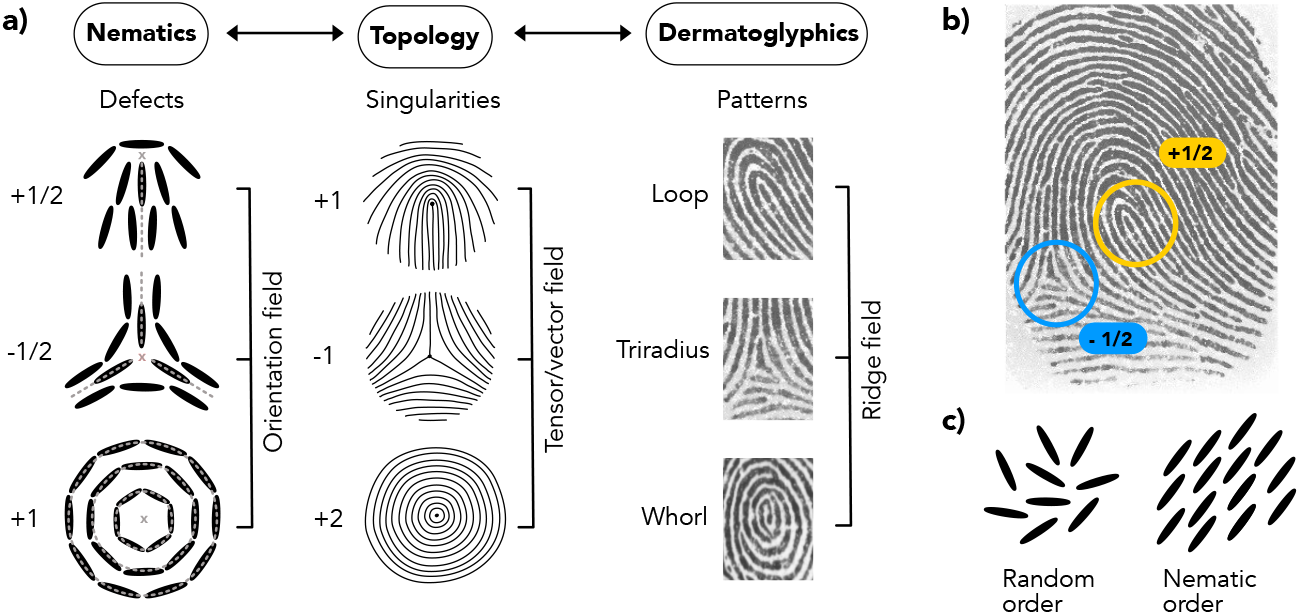
Parallels between dermatoglyphics, topology, and nematics. (a) Comparison of concepts and properties across the three fields, highlighting the correspondence between nematic defects, topological singularities, and dermatoglyphic patterns. (b) Example fingerprint showing 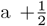 defect (yellow) and 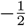 defect (blue), giving a total topological charge of zero. (c) Arrangements of units in random and nematic order. Winding of nematic defects is shown in Supplementary Section S6. Fingerprint images adapted from the Fingerprint Verification Competition (FVC) databases [11, 12].

In this work, the ridge network is treated as a tangential director line field defined on the surface of the fingertip. This field encodes ridge orientation but not polarity, consistent with the head–tail symmetry of nematic director fields [2]. The director field description allows ridge orientation to be treated as a continuous field, independently of ridge spacing or thickness.

Penrose relates loops and triradii to digit number using a global topological tensor field [9]. In this framework, whorls are treated as two loops, and the analysis applies at the scale of the entire hand rather than the individual fingertip. By contrast, the present work treats each fingerprint as a compact, bounded domain, with defect configurations dictated by the balance between boundary and interior winding. Analogies between nematic liquid crystals and fingerprints have also been demonstrated using similar image analysis techniques [13]. Furthermore, the extensive study of nematics on curved surfaces and complex geometries provides ready-made theorems that can be applied to the fingerprint.

## 4 A mathematical nematic approach

The fingerprint ridge network is confined to a bounded patch of the fingertip surface, whose functional area is scanned for biometric identification. This region is bounded superiorly by the nail edge, laterally by the flattened sides of the finger, and inferiorly by the proximal crease. Topologically, this domain is homeomorphic to a two-dimensional disk. Such compact, connected surfaces are characterised by an Euler characteristic *χ*(*S*), which for a simple bounded disk is equal to one [14].

The Poincaré–Hopf theorem relates the total index of singularities in a vector field to the Euler characteristic of the underlying manifold [14, 15]. For nematic director fields, which are invariant under 180° rotation, the Poincaré–Hopf theorem generalises to allow half-integer defects through a projective formulation [2, 15]. For compact surfaces with boundaries, the topological constraint on total defect charge is modified by boundary conditions. The boundary winding number *W*_∂*S*_ describes the net rotation of the director field along the boundary of the domain [16]. For a nematic director field on a bounded surface, the sum of defect charges and the boundary winding together satisfies

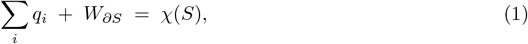

where ∑_*i*_ *q*_*i*_ is the sum of interior defect charges [15]. This approach is shown in Fig. 2. A detailed derivation of this relation for nematic director fields on bounded surfaces, including intermediate steps, is provided in the Supplementary Information (Sections S2–S5). For idealised fingerprint geometry, the boundary winding is close to one, consequently, the sum of defect charges is predicted to be zero (Supplementary Sections S4–S5).

**Figure 2.**
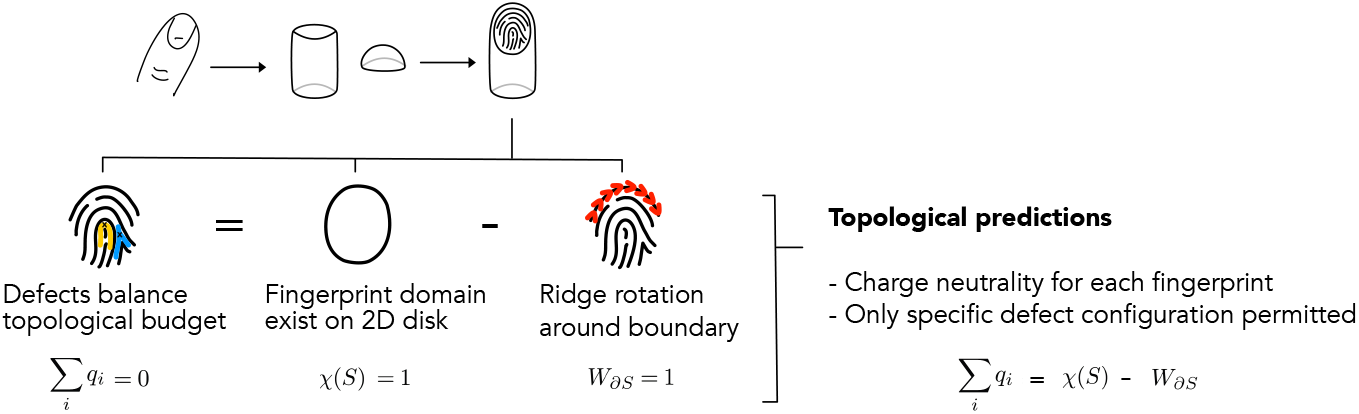
Geometric and topological prediction of fingerprint charge neutrality. The fingertip is modelled as a cylinder–hemisphere composite, with the fingerprint domain forming a disk-like region of Euler characteristic *χ*(*S*) = 1. In the idealised case, ridge orientation rotates by 180^*°*^ around the domain boundary (red arrows), giving nematic boundary winding *W*_∂*S*_ = 1. Applying the Poincaré–Hopf theorem to an illustrative fingerprint containing one loop (+1*/*2, yellow) and one triradius ( 1*/*2, blue) yields a total defect charge of zero: ∑_*i*_ *q*_*i*_ = *χ*(*S*) *W*_∂*S*_ = 1 1 = 0. This theorem therefore imposes a charge-neutrality constraint on each fingerprint domain, permitting only certain defect configurations.

## 5 Empirical validation using fingerprints

To test the predicted topological constraint, 133 fingerprints were analysed from open-access databases curated for fingerprint recognition research [11, 12], retaining one complete impression per finger. Defects were identified by visual inspection of ridge orientation patterns following established dermatoglyphic conventions [17, 18], with triradii, loops, and whorls classified as 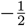, 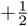, and +1 defects respectively [2, 9] (Fig. 1).

Visual classification remains the basis of fingerprint analysis and is robust for identifying ridge singularities [18]. Equivalent approaches are used in nematics, where singularities are defined by director winding rather than microscopic structure [13]. Across the dataset, 127 out of 133 fingerprints (95.5%) satisfied the predicted neutrality condition, with total defect charge summing to zero (Fig. 3a).

**Figure 3.**
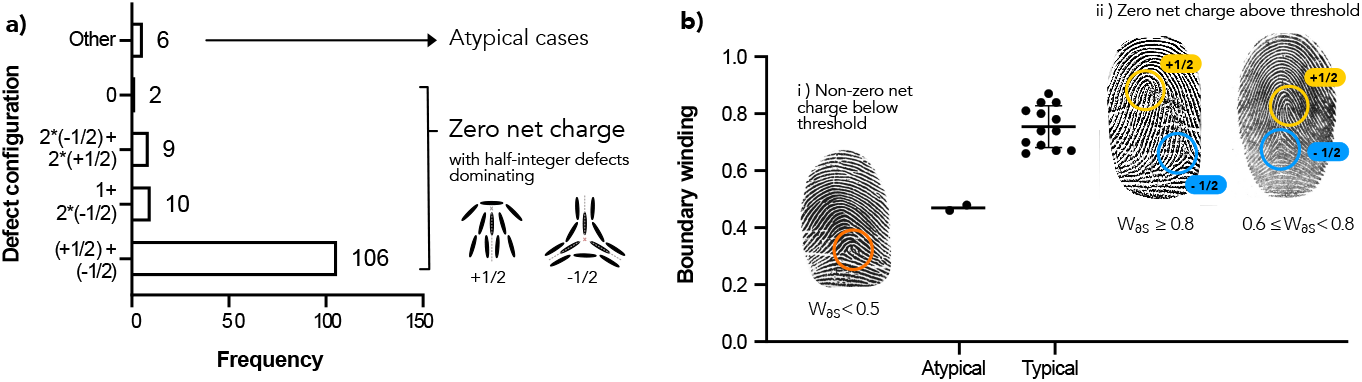
Frequency distribution of fingerprint configurations and boundary winding. (a) Frequency of defect configurations across the dataset: 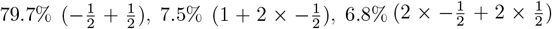, 1.5% (0), and 4.5% (other). Overall, 95.5% of fingerprints satisfy the zero net topological charge prediction, dominated by configurations containing one of each half-integer defect (79.7%, *n* = 106). Atypical configurations (4.5%, *n* = 6) do not exhibit neutrality. (b) Boundary winding analysis (*n* = 15). Typical fingerprints satisfying charge neutrality cluster between 0.66–0.87 (87%). Representative examples spanning this range are shown; 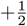 defects are labelled in yellow and 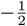 in blue. Atypical boundary winding occurs below 0.5 (13%); an example is shown with a difficult-to-define defect highlighted in orange. Fingerprint images adapted from FVC databases [11, 12].

The dominant configuration (79.7%) comprised one 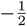 defect paired with one 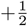 defect. Less frequent configurations also conserved zero net charge, including two 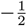 defects with one +1 defect, and two 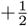 defects with two 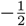 defects. The observed distribution is similar to that reported in larger dermatoglyphic studies [17].

Defect configurations deviated significantly from uniformity (*χ*^2^ = 297.71, *p* = 3.37 × 10^−63^), while global charge was tightly conserved (median = 0; 95% bootstrap CI = [0, 0]; mean = −0.008 ± 0.106 SD). A small fraction (4.5%) did not satisfy global neutrality; these atypical prints contained ridge configurations that were difficult to assign to standard (half-)integer charges (Fig. 3) and were assigned the closest half-integer consistent with ridge geometry. Boundary winding was measured for a subset (*n* = 15; 13 charge-neutral and 2 atypical) by tracking the net rotation of ridge orientation along the fingerprint boundary. Mean boundary winding was *W*_∂*S*_ = 0.71 *±* 0.12 (range 0.46–0.87), lower than the idealised *W*_∂*S*_ = 1 (Fig. 3b). The subset showed a clear threshold structure: fingerprints with *W*_∂*S*_ ≥ 0.6 were charge-neutral, whereas atypical cases occurred at *W*_∂*S*_ *<* 0.5 and exhibited non-zero net charge. Boundary-induced deviations and charge-quantisation effects are discussed in Supplementary Sections S7–S8.

## 6 Discussion

Existing models of fingerprint formation focus primarily on generative mechanisms, including biochemical signalling and mechanical instabilities that initiate and propagate ridges during development [8, 19]. While these approaches successfully explain how fingerprint ridges arise, they do not consider the restrictions that permit specific configurations. By contrast, the nematic framework identifies a global topological constraint that fingerprint patterns must satisfy, independent of the specific biological processes involved. The inherently stochastic reaction-diffusion system responsible for biochemical patterning of ridges [8] provides the uniqueness of fingerprint patterns, while the topology and boundary conditions constrain the configurations that emerge. The empirical finding that average *W*_∂*S*_ = 0.71, rather than the idealised *W*_∂*S*_ = 1, does not undermine the framework; it exposes a physical limitation. Because defect charges are discrete (half-integer quantised) whereas boundary winding varies continuously, exact cancellation is not always possible. Nevertheless, fingerprints with *W*_∂*S*_ ≥ 0.6 satisfy charge neutrality, whereas those below this threshold show geometric frustration and atypical defect configurations. This behaviour plausibly reflects finite ridge width and discrete defect structure, which coarse-grain the idealised continuous director field. The topological constraint therefore remains valid, but its physical realisation is moderated by material constraints in fingerprints.

In this view, fingerprints are a passive biological record of orientational order on a confined surface. Once formed, ridge patterns persist unchanged, preserving the topological structure set during development. Variation in boundary geometry alters boundary winding and thus the defect configurations observed across fingerprints. The threshold at *W*_∂*S*_ ≈ 0.6 reflects the interplay between continuous topology (which would permit arbitrary *W*_∂*S*_) and discrete physical realisation, which coarse-grains ∑_*i*_ *q*_*i*_. This offers a geometric explanation for population-level variability without requiring differences in developmental pathways.

Related links between topology, geometry, and defect organisation occur across biological nematic systems, including epithelial tissues and cytoskeletal assemblies [1–7, 20]. Fingerprints therefore provide an accessible macroscopic example of principles that operate across scales in living matter. We do not model the dynamics of ridge formation, nor do we suggest that fingerprints behave as active nematics during development. Rather, we show that final ridge configurations are subject to global physical constraints.

## 7 Limitations

Defect identification relied on visual classification of ridge patterns, which may introduce subjectivity, particularly in atypical cases or near boundaries. The fingerprints analysed were drawn from biometric databases rather than controlled developmental studies, limiting access to information about age, genetic background, or developmental conditions. These limitations do not affect the topological nature of the analysis, which depends only on ridge orientation and boundary geometry rather than on developmental variables.

## 8 Conclusion

Fingerprint ridge patterns can be understood as frozen nematic fields defined on a bounded surface. Their global organisation is constrained by topology and boundary geometry, leading to a conserved total defect charge across individuals. This framework explains both the diversity and regularity of fingerprint patterns and demonstrates that global physical constraints play a fundamental role in biological pattern formation.

## Acknowledgements

Supported by the ARIA “Nature Computes Better” Opportunity Space, (Bio)active Matter Based Computation. K.W conceived the project, supervised and obtained funding. L.M. originated the fingerprint concept and led data analysis. L.M. wrote the original draft and K.W. revised the manuscript. We acknowledge the FVC consortium for making fingerprint databases publicly available. In memory of Edith Merrett.

