## Supplementary Information for "Fingerprints as Frozen Nematic Fields"

### SI–Theory: Topological Framework for Fingerprints

#### S1. Nematic director fields and defect charge

Fingerprint ridge orientation is treated as a two-dimensional nematic director field with head–tail symmetry ($\mathbf{n} \equiv -\mathbf{n}$). Singularities correspond to topological defects, whose charge $q$ is defined by the net winding of the director field around a closed loop.

Because nematic director orientation is defined modulo $\pi$, defects with half-integer charge are permitted. In fingerprints, triradii, loops, and whorls correspond to $-\frac{1}{2}$, $+\frac{1}{2}$, and $+1$ defects respectively.

#### S2. Vector and nematic formulations of the Poincaré–Hopf theorem

The classical Poincaré–Hopf theorem states that the sum of indices of isolated singularities in a vector field equals the Euler characteristic $\chi(S)$ of the surface:

$$\sum_i q_i = \chi(S). \tag{1}$$

For nematic director fields, which are invariant under 180° rotation, the theorem generalises to a projective formulation admitting half-integer indices, with defect charges $q_i$ constrained by surface topology.

#### S3. Surfaces with boundary and boundary winding

For compact surfaces with boundary, the Poincaré–Hopf relation includes a boundary contribution. For nematic director fields on a bounded surface $S$:

$$\sum_i q_i + W_{\partial S} = \chi(S), \tag{2}$$

where $W_{\partial S}$ is the boundary winding number, defined as the net rotation of the nematic director field along the boundary $\partial S$.

Boundary winding captures the geometric contribution of the boundary to total topological charge and varies continuously with boundary geometry.

#### S4. Topology of the fingerprint domain

The fingerprint ridge field is defined on the functional contact area of the fingertip, which is compact, connected, and bounded by a single closed curve, and therefore topologically equivalent to a disk. :

$$\chi(S) = 1. \tag{3}$$

Substituting Eq. 3 into Eq. 2 yields:

$$\sum_i q_i + W_{\partial S} = 1. \tag{4}$$

#### S5. Boundary winding in fingerprints and the neutrality prediction

For an idealised fingerprint geometry, ridge orientation varies smoothly along the boundary below the nail. Traversing the boundary once, ridges rotate by approximately 180° from the lateral sides to the superior nail edge. Near the proximal crease of the idealised disk domain, ridges do not remain tangent to the boundary; instead, the initiation site above the crease interrupts orientation, so the director field does not complete a full 360° rotation. For an idealised fingerprint with nematic symmetry, boundary winding is close to unity:

$$W_{\partial S} \approx 1. \tag{5}$$

Substituting Eq. 5 into Eq. 4 gives the neutrality prediction:

$$\sum_i q_i \approx 0. \tag{6}$$

Thus, the total nematic defect charge within a fingerprint is predicted to be zero.

#### S6. Discrete defect charges and permitted configurations

Nematic defect charges are quantised in half-integer units. In fingerprints, the neutrality condition in Eq. 6 can therefore be satisfied by multiple combinations of triradii, loops, and whorls. For example, one $-\frac{1}{2}$ defect paired with one $+\frac{1}{2}$ defect satisfies the constraint, as do configurations involving integer defects provided the total charge sums to zero. Quantisation restricts admissible configurations while still allowing substantial variability in pattern type.

#### Winding of (half-)integer nematic defects

The topological charge $q_i$ of a nematic defect is set by the net director rotation around a loop enclosing the defect centre (singularity):

$$q_i = \Delta\theta/\pi \tag{7}$$

where $\Delta\theta$ is the total change in director orientation (radians) over one circuit. Because nematic directors are defined modulo $\pi$, half-integer defect charges are permitted.

##### -1/2 defect (triradius)

In fingerprints, a triradius occurs where three ridge fronts meet at approximately 120° angles. Traversing a closed loop around the defect, the director rotates clockwise by 90°, giving

$$q_i = \Delta\theta/\pi = (-\pi/2)/\pi = -1/2 \tag{8}$$

This yields the triradius pattern as ridges converge from three directions while the director rotates opposite to the traversal direction.

##### +1/2 defect (loop)

A loop forms where ridges curve back on themselves, creating a U-shaped pattern. Around a closed loop enclosing the defect, the director rotates by 90° in the same direction as the traversal, giving

$$q_i = \Delta\theta/\pi = (+\pi/2)/\pi = +1/2 \tag{9}$$

The positive charge corresponds to director rotation aligned with the traversal direction, producing the characteristic loop geometry.

##### +1 defect (whorl)

A whorl exhibits concentric circular ridges, with the director forming closed streamlines around the centre. This yields

$$q_i = \Delta\theta/\pi = \pi/\pi = +1 \tag{10}$$

Here the director completes one full nematic rotation about the defect centre (equivalent to 180° under $\mathbf{n} \equiv -\mathbf{n}$).

#### S7. Boundary winding and physical realisation

The interaction between continuous boundary geometry and discrete defect charges imposes a physical constraint on fingerprint configurations. Boundary winding $W_{\partial S}$ varies continuously with boundary geometry, whereas defect charges are quantised in half-integer units, so exact cancellation is not always possible. For idealised geometry with $W_{\partial S} = 1$, Eq. 2 predicts $\sum_i q_i = 0$. Empirically, $W_{\partial S} = 0.71 \pm 0.12$ (mean $\pm$ SD), which would formally predict $\sum_i q_i \approx +0.29$. Because available defect charges are restricted to a discrete set $(\ldots, -\frac{1}{2}, 0, +\frac{1}{2}, +1, \ldots)$, the system must select the closest realisable configuration.

Applying Eq. (1) suggests that when boundary winding falls below $W_{\partial S} \approx 0.5$, discrete defect charges and finite ridge width produce a mismatch between interior charge and boundary winding. In this regime, the predicted interior charge deficit ($\sum_i q_i = +0.5$ to $+0.6$) may exceed what the ridge network can physically accommodate, leading to non-neutral configurations.

Finite ridge width, non-ideal fingertip geometry, and quantised defect charges may impose a coarse-graining on realisable configurations. In a subset of 15 fingerprints (13 typical and 2 atypical), neutrality was maintained for $W_{\partial S} \geq 0.6$ (13/15), consistent with selection of $\sum_i q_i = 0$ as the closest admissible configuration. For $W_{\partial S} < 0.5$, fingerprints showed non-neutral configurations, including defects that cannot be assigned to conventional nematic charges, consistent with the predicted interior charge deficit ($\sum_i q_i = +0.5$ to $+0.6$) becoming physically unrealisable.

#### S8. Boundary winding threshold

The empirical boundary-winding threshold ($W_{\partial S} \approx 0.6$) may mark a regime below which geometric frustration prevents charge cancellation. Rather than violating Eq. 2, these cases illustrate how a continuum topological prediction is realised in a discrete biological system. The topological relation remains valid, while material constraints modulate its physical realisation across boundary geometries.

These atypical cases reflect boundary geometry and physical realisation rather than violations of the underlying topological constraint, illustrating how a continuum topological prediction is realised in a discrete biological system.

#### S9. Relation to prior topological descriptions of fingerprints

Previous topological descriptions treated the hand as a global object, relating pattern counts to digit number. Here, each fingerprint is treated as an independent bounded domain and nematic defect theory is applied locally. This formulation naturally incorporates half-integer defects and boundary contributions, providing a unified framework for fingerprint pattern constraints.

### SI–Methods: Empirical Fingerprint Analysis

To validate the mathematical prediction of zero net defect charge, we analysed real human fingerprint samples.

#### Fingerprint defect analysis

We collected 133 fingerprint samples from open-access databases originally curated for fingerprint recognition research. To avoid duplication, only one sample per finger was retained. For each fingerprint, visible ridge defects were identified and classified by observation using the dermatoglyphic–nematic equivalencies. Defects were counted by visual inspection of the ridge orientation field by one observer and verified against characteristic nematic configurations. The total number of each defect type and the resulting net topological charge were then calculated.

#### Boundary winding analysis

To assess variation in boundary geometry, ridge orientations were measured at 8 evenly spaced points around the boundary for a subset of 15 fingerprints (chosen to include 13 typical charge-neutral and 2 atypical cases to represent the wider dataset) using ImageJ. The boundary winding number $W_{\partial S}$ was calculated as

$$W_{\partial S} = \frac{\theta_{\max} - \theta_{\min}}{180}, \tag{11}$$

where $\theta_{\max}$ and $\theta_{\min}$ are the maximum and minimum ridge angles (degrees). This metric quantifies the total angular excursion of ridges around the boundary, with $W_{\partial S} = 1$ corresponding to the idealised 180° rotation expected for a perfect disk geometry.

#### Statistical analysis

To test whether the observed distribution of defect configurations deviated from uniformity, a chi-squared goodness-of-fit test was performed using Python. The null hypothesis assumed equal frequency across all possible defect configurations. The test statistic was calculated as

$$\chi^2 = \sum \frac{(Observed - Expected)^2}{Expected}, \tag{12}$$

across five configuration categories: $(-\frac{1}{2} + \frac{1}{2})$, $(1 + 2 \times -\frac{1}{2})$, $(2 \times -\frac{1}{2} + 2 \times \frac{1}{2})$, $(0)$, and other.

With 133 fingerprints across five categories, the test yielded $\chi^2 = 297.71, p = 3.37 \times 10^{-63}$. This significant result demonstrates that fingerprint configurations are highly non-uniform, with a strong preference for charge-neutral patterns, particularly the single $(-\frac{1}{2} + \frac{1}{2})$ pairing (79.7% of cases).

To assess robustness of charge conservation, a 95% bootstrap confidence interval for the median total defect charge was calculated using Python. Using 10,000 bootstrap resamples with replacement from the 133-fingerprint dataset, median charge values were computed for each resample. The resulting 95% confidence interval was $[0, 0]$, with both the 2.5th and 97.5th percentiles equal to zero. This tight interval reflects the strong tendency towards charge neutrality and demonstrates the robustness of the observed charge conservation.
